# A country-wide examination of effects of urbanization on common birds

**DOI:** 10.1101/2022.11.27.518114

**Authors:** Lyanne Brouwer, Lisenka de Vries, Henk Sierdsema, Henk van der Jeugd

## Abstract

Urbanization forms one of the most drastic alterations of the environment and poses a major threat to wildlife. The human-induced modifications of the landscape may affect individual’s fitness and thereby result in population declines. Research on how urbanization affects fitness traits has shown mixed results, but typically contrasted data from few urban and non-urban sites collected over short time frames from single species. It thus remains unknown whether we can generalize across species, whereas such knowledge is crucial for population predictions that are needed for conservation management. Here, we use data from a nation-wide citizen science project to examine variation in survival and body mass and size of common passerine birds, collected along an urbanization gradient in the Netherlands over an 8-year period. Although the overall association between urbanization and survival was slightly negative, there was strong support for lower survival in three species, and higher survival in two of the 11 species examined. Effects of urbanization on body mass and size also varied but were far less strong and there was no evidence that they mediated the impacts on survival. Our results imply that body mass and size cannot be used as indicators for urban-associated patterns of survival. Furthermore, the species-specific survival responses indicate that care should be taken when predicting the effects of ongoing urbanization for communities, because even closely related species can show different responses. Moreover, the contrasting survival successes among species suggests that ongoing urbanization may lead to shifts in community structure and loss of biodiversity.

## Introduction

In Europe the expansion of urban land use (‘urban sprawl’) has increased by 5% between 2006-2009 (European Environment Agency, 2016). Urbanization involves some of the most drastic alterations of the environment through increased habitat fragmentation (Crooks, Suarez, & Bolger, 2004; Fischer & Lindenmayer, 2007), higher disturbance from humans (Fernandez, Lopez-Calleja, & Bozinovic, 2002), introduction of novel predator communities (Møller & Ibáñez-Álamo, 2012; Bonnington, Gaston, & Evans, 2013), increased light at night-time (Spoelstra & Visser, 2013; Aulsebrook et al., 2020) and noise (Halfwerk et al., 2011; Potvin, Mulder, & Parris, 2014). These anthropogenic disturbances can have large implications on biodiversity as they may change an organism’s fitness, which in turn may lead to population declines. Despite these challenges, cities and towns have become important habitats by supporting a significant proportion of the world’s biodiversity (Mcdonald, Kareiva, & Forman, 2008; Aronson et al., 2014). In fact, some species even thrive in urbanized areas (McKinney, 2006; Isaksson, 2018). Thus, the human-built environment offers a range of challenges and opportunities, and the predicted ongoing urbanization (Gao & O’Neill, 2020) means that it will be of utmost importance for conservation management to understand the consequences this will have on wildlife populations.

Survival is a major fitness component and understanding how survival of wildlife responds to urbanization will increase knowledge on the processes that regulate species abundance in such environments. For example, although some species may occur in cities, low survival could indicate that these populations are not self-sustained but driven by dispersal of juveniles into cities (Withey & Marzluff, 2005). Survival of birds in urban areas may be low because they suffer immediate lethal consequences from fatal collisions with buildings (Elmore et al., 2020) or from feral predators, which occur at high density in urban areas (Loss, Will, & Marra, 2013; Legge et al., 2017; but see Fischer et al., 2012). On the other hand, higher year-round resource availability resulting from the extended plant growing seasons (Jochner et al., 2013) and the availability of anthropogenic food sources may allow some species to increase their survival prospects.

A recent meta-analysis across ten studies on 15 bird species found higher survival in response to urbanization, which was suggested to be one of the most convincing intraspecific trends in life-history traits observed along the urbanization gradient (Sepp et al., 2018). However, given that species do not uniformly suffer or benefit (McKinney, 2006; Isaksson, 2018), it is unclear whether such a generalization across all species is meaningful. Furthermore, variation in how urbanization affects survival (e.g. Marzluff & Neatherlin, 2006; Horak & Lebreton, 2008; Evans et al., 2015), may also reflect geographic variation or be the result from specific study designs. Most studies compared survival among the extreme ends of the urbanization gradient (notably urban versus rural populations but see: Evans et al., 2015). However, this ignores intermediate habitats such as suburban areas that might be particularly relevant, because they typically cover significant areas and hold high populations of bird species (Cannon, 1999). So far, large scale multi-species investigations of survival along a broad urban gradient is limited to a study from north-eastern USA, which found mixed survival responses with particularly the more generalist species benefitting from urbanization (Evans et al., 2015). Whether this is a general pattern remains to be investigated.

We have even less knowledge about how urbanization alters physiological and behavioural processes that may result in variation in body mass and size of individuals. Since adult body mass reflects the degree to which an individual has fat reserves (Labocha & Hayes, 2012), mass may directly impact reproduction and survival (Verhulst et al., 2004; Cresswell, 2009) and could thus reveal more details on the underlying mechanisms that regulate populations. In addition, if urbanization affects mass and survival in a similar way (or in opposite directions), the former may serve as an indicator that does not require multiple years of data collection. However, creating predictions for the response of body mass to urbanization is not straightforward. Positive associations can be expected because the high predicted food availability in urban areas will enable individuals to carry sufficient fat reserves that can serve as a buffer against potential food shortage in the future (Cresswell, 2009). On the other hand, negative associations can be predicted because fat reserves also come with costs to locomotion and metabolism (Witter & Cuthill, 1993). Thus, the highly predictable continuous input of food in urban areas will result in wildlife being less dependent on fat reserves and live on the credit of tomorrow’s food (“credit card hypothesis”, Shochat, 2004).

Anthropogenic food sources may be of insufficient quality and negatively affect development and growth, particularly during the nestling or juvenile stage (Seress et al., 2020). At the same time, our cities are urban heat islands which experience higher temperatures than the surrounding areas (Merckx et al., 2018). For example, temperatures in the Amsterdam region of the Netherlands were shown to be over 3^#x25A1;^C higher compared to the surrounding countryside on moderately warm days (Koomen & Diogo, 2017). Warming temperatures have been suggested to select for smaller body sizes (Sepp et al., 2018), because higher ambient temperatures increase metabolic rates and the associated costs for a given body size (Brown et al., 2004). Thus, both lower food quality and urban heat island effects mean that urbanization is expected to be associated with smaller size.

The reported associations between urbanization and body mass and size show mixed results (for review see: Sepp et al., 2018 and references therein). However, it should be noted that most studies are based on a single snapshot in time and compare small numbers of ‘urban’ and ‘rural’ locations. This may be problematic since a study comparing body mass and size of blackbirds (*Turdus merula*) among 11 paired ‘urban’ and ‘rural’ sites showed that the magnitude and direction of responses varied among sites (Evans et al., 2009). Large-scale studies examining survival and biometry of multiple species along an urbanization gradient are thus crucial to better understand the differences in urbanization effects across species.

Here, we investigate effects of urbanization on survival and biometry in 14 common passerine bird species (see Table S1), to determine whether there are general patterns in how fitness traits of such species respond to urbanization. Data was collected through capture-mark-recapture in a citizen science project along an urban gradient (>200 locations) in the Netherlands. This densely populated Western European country has a long history of urbanization, and the urban area now covers 16% of the country’s surface, which forms an important part of the breeding habitat for many bird species (Snep et al., 2015). To capture variation in land-use within urban areas (e.g. parks, buildings), we have not only defined urbanization as the distance from the city’s border, but additionally used the proportion of area covered by impervious surface (IMP, roads and buildings). Since larger individuals are also heavier, but do not necessarily carry more body fat, we use structural equation modelling to determine the relative importance of urbanization on mass and size, while accounting for the correlation between mass and size. According to the urban heat island effect, we predict smaller size with increased urbanization. In addition, the lower dependency on fat reserves in urban areas is expected to result in lower body mass, which is predicted to be particularly pronounced for generalist species that are expected to be able to benefit from human derived food most.

## Methods

### Data collection

Data utilized here combines data from >84,000 individuals of 14 common bird species which were encountered along the urban gradient (Table S1). Data was obtained through capture-mark-recapture from three citizen science projects carried out from 2011 to 2019 at 217 locations throughout the Netherlands (Fig. 1a). Each of the projects followed specific standardized procedures. First, the Dutch constant effort site (CES) project follows European protocols and collects capture-mark-recapture data 12 times per breeding season (13 April-13 August) for long-term monitoring of bird populations in rural areas (Robinson, Julliard, & Saracco, 2009). Second, in the ‘*ring*-MUS’ project, a citizen science project coordinated by the Dutch Centre for Avian Migration and Demography with support from BirdLife Netherlands, mark-recapture data were collected at least twice a month at private residences within the urban and suburban matrix, providing the data in built-up areas which are not normally collected in the CES project. Third, for some species (common starling (*Sturnus vulgaris*), house sparrow (*Passer domesticus)*, common blackbird (*Turdus merula*), European greenfinch (*Chloris chloris*), great tit (*Parus major*) and blue tit (*Cyanistes caeruleus*)) additional data was available through species specific projects (RAS projects: Re-trapping Adults for Survival). In such projects the focus was on capturing all individuals of a single species in a given area, from varying habitat types along the urban gradient. In addition to mist netting, catching techniques involved walk-in traps, clap traps and catching birds in nest boxes. To reduce bias with respect to differences in timing of data collection among the projects, the same recapture period was selected from each of the three projects (i.e. the CES period: 13 April-13 August), thereby also specifically focussing on resident birds on their breeding grounds.

**Fig 1a).**
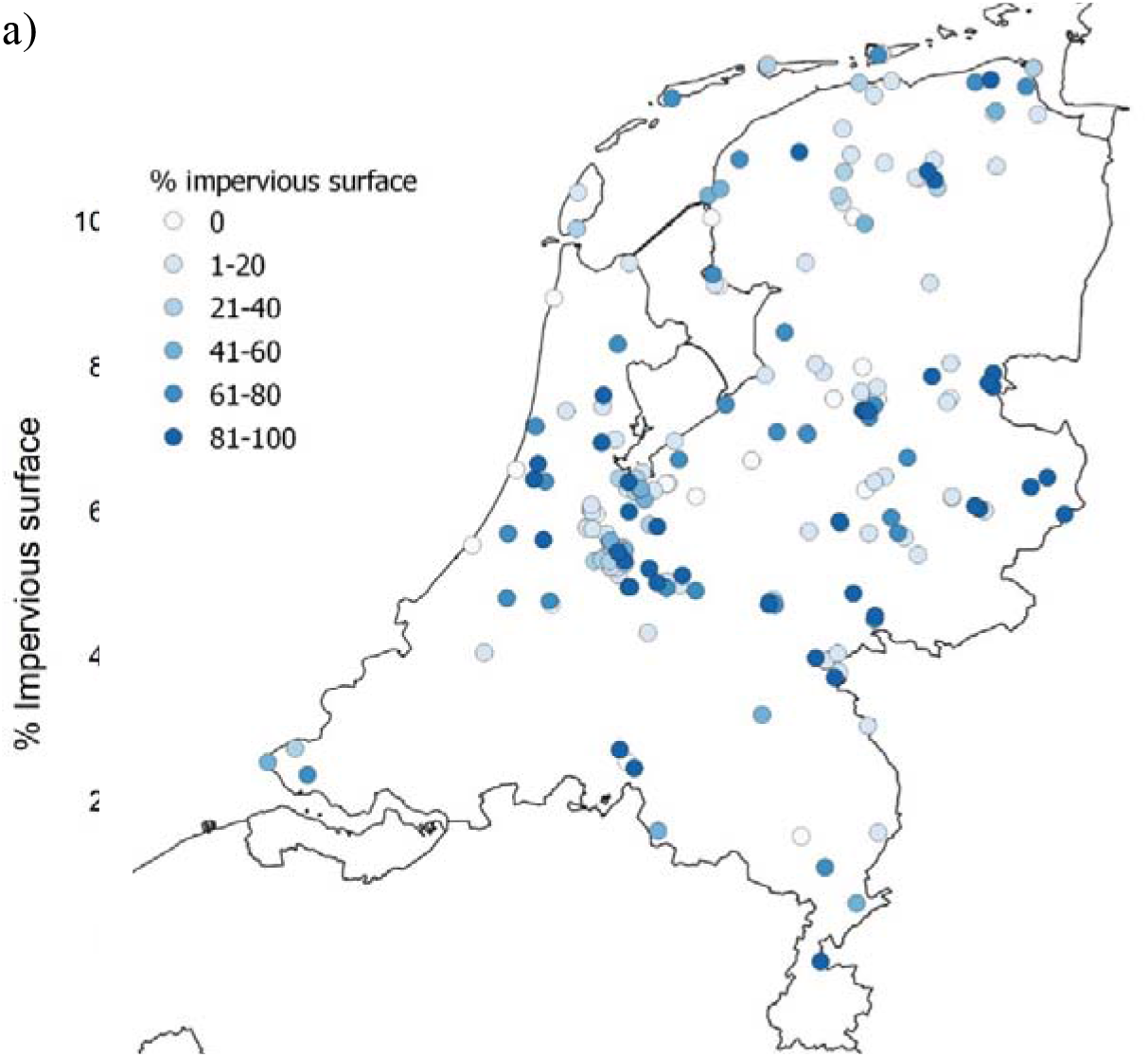
Map showing capture locations with different colours indicating the degree of urbanization measured as the percentage impervious surface. b) The association between percentage impervious surface and distance from the city border (with negative values indicating rural and positive urban areas) for each of the 217 capture locations.

All birds were banded with uniquely numbered alloy split bands, allowing for individual identification when recaptured at a later occasion. In addition, in both *ring*-MUS and RAS projects some species (house sparrow, common blackbird, common starling, European greenfinch and great tit) were colour banded, allowing for identification of individual birds through resightings. Data was collected by licensed volunteers, who received training in handling, banding and processing birds using standardized methods. Captured birds were aged and sexed based on plumage characteristics where possible, and when time allowed morphometric measurements were taken, including body mass to the nearest 0.1g with an electronic balance and wing length (maximum flattened chord length) to the nearest 0.5mm using a butt-ended ruler.

### Degree of urbanization

For each of the 217 capture/resighting locations the percentage impervious surface (IMP) within a 200m radius (size area=12.6 ha) and the distance from the city border (with negative values indicating rural and positive urban areas) were calculated using software ArcGIS (Environmental Systems Research Institute, 2011). GIS data on land use including city borders were obtained from Statistics Netherlands (https://www.pdok.nl/introductie/-/article/cbs-bestand-bodemgebruik, version 2015). From these we selected infrastructure, residential areas (including gardens, of which most of their surface is paved (Stobbelaar, van der Knaap, & Spijker, 2021)), retail areas and industrial areas to calculate IMP. Distance from the city border was highly correlated with IMP (*r*=0.69), nevertheless there was still considerable variation in IMP within urban areas (Fig. 1b).

### Survival analyses

To estimate whether annual adult survival is associated with urbanization, capture-recapture histories were created for each banded individual, with a ‘1’ if the bird was captured or resighted any time within a breeding season and a ‘0’ otherwise. Using the constructed capture histories apparent annual survival between breeding seasons was calculated with a live-recapture model in RMark (Laake, 2013). To account for variation in survival and recapture/resighting rates among projects and groups of individuals (e.g. ring type), *a priori* models were constructed for each species separately as follows: recapture rates were allowed to vary among projects (CES, *ring*-MUS, RAS), ring type (colour or metal only), age class (first year versus older individuals), age at first capture (ring-age) and their interactions; survival was allowed to vary among age classes, although we only examined effects of urbanization on adult survival, since the dispersive nature of juveniles means that their survival is a lot harder to estimate accurately. To avoid over-parametrization, we did not fit full time-dependent models. Models were further simplified where possible (i.e. not all species were colour banded or part of the RAS project) to derive a baseline (null) model which could be used as a starting point for our analyses (see Table S1 for details). We then tested whether adult survival was associated with urbanization by including the distance from the city border and IMP as covariates for the adult survival parameter. In addition, to investigate any non-linear association between adult survival and urbanization, models including the quadratic term of these covariates were also run. The urbanization covariates were scaled to z-scores to facilitate model convergence.

Previous work has indicated the presence of resident and non-resident (migratory, i.e. transient) individuals in the dataset (Johnston et al., 2016). Such transients have a low probability of being reencountered, violating the capture-recapture assumption that all individuals have equal recapture probabilities (Lebreton et al., 1992). To account for transients, an extra time step after the first occasion was added to the capture history (Johnston et al., 2016). Individuals that have been captured more than once during the first capture season are assumed to be residents. For these individuals, a ‘1’is inserted into the capture history after the first occasion. Birds that have been captured only once can either be a resident or a transient individual. For these individuals, a ‘0’ is inserted into the capture history after the first occasion. Using this approach allows for the separate estimation of a ‘transience probability’ with the estimated ‘Phi’ being the probability of a bird being a resident, while the estimated recapture gives the probability that a resident bird is identified as such (Johnston et al., 2016). Effectively, this means individuals that have only been caught once do not contribute to the estimation of the survival probability (Pradel et al., 1997). Unfortunately, using this method does not allow for goodness-of-fit testing.

Low recapture rates and/or ringing effort resulted in non-identifiable parameters in survival models for song thrush (*Turdus philomelos)*, common chaffinch (*Fringilla coelebs*) and great spotted woodpecker (*Dendrocopos major*) and therefore only effects on biometry were considered for these species. For the remaining 11 species ringing effort varied between 2,300-14,582 per species (Table S1).

Model selection was based on Akaike’s Information Criterion corrected for sample size (AICc; Akaike, 1973), with lower AICc values being considered as better supported by the data. In addition, we report normalized Akaike weights to assess the relative support for competing models (Burnham & Anderson, 2002).

### Biometry analyses

Since we aimed to investigate how both adult body mass and size vary with urbanization, but mass is also affected by size, we constructed structural equation models using R package piecewiseSEM (Lefcheck, 2016). These models allow for the evaluation of causal linkages among variables in a single multivariate framework. Wing length was used as an indication of size, whereby both body mass and wing length were fitted using a Gaussian distribution, with either distance from the city border or IMP as predictors (either linearly or quadratic). To account for the fact that larger individuals are also heavier, wing length was included as a linear predictor for body mass too (see Fig. S1 for path diagram), thereby effectively analysing relative body mass. We also accounted for potential confounding variables by including project and sex as fixed factors, and Julian day as a covariate in the body mass and wing length equations. In addition, to account for variation in mass during the day, time of weighing since sunrise was calculated using package StreamMetabolism (Sefick Jr., 2016). All variables were scaled to z-scores before including them in the model. Individual identity, year and identity of the capture location were included as random intercepts to account for non-independence of the data. For great spotted woodpecker, common blackbird, common starling and Eurasian wren (*Troglodytes troglodytes*), the random intercept for year led to convergence issues and was therefore omitted from the model. Limited data on biometry measurements meant that effects on mass and size could not be examined for European robins (*Erithacus rubecula*). We evaluated the models based on the global goodness of fit of the null model (without urbanization predictors), which showed that the models fitted the data well (Fisher’s C test, all *P*>0.07). The importance of the urbanization predictors for explaining variation in relative mass and size was based on their estimated effect sizes and 95% CI’s combined with the marginal (proportion of total variance explained by the fixed effects) and conditional (proportion of total variance explained by both fixed and random effects) *R*^2^ (Nakagawa, Johnson, & Schielzeth, 2017).

## Results

### Survival

The mark-recapture models showed that after accounting for variation in recapture rates (e.g. among the different projects), there was strong variation in both magnitude and direction of the effects of urbanization on apparent adult survival between breeding seasons (Fig. 2a). There was little evidence for a general pattern with respect to effects of urbanization on survival across species: despite the weighed mean effect sizes of the association between survival and urbanization being negative, the 95% CI’s overlapped zero (Fig. 2a). Model selection results showed that at least one of the two urbanization parameters used as a predictor for variation in survival was strongly supported by the data in five of the 11 examined species (Table 1). Three species (chiffchaff, European robin, European greenfinch) showed reduced survival closer to the city centre (Fig. 2a) and this reduction was substantial, with respectively ∼50%, ∼25% and ∼13% lower survival for greenfinches, robins and chiffchaffs living in city centres compared to those living 3.5km from cities (Fig. S2). Although the estimated effect sizes for common starlings suggested a strong reduction in survival with increased urbanization, the CI’s were large (Fig 2a) and models including quadratic effects did not converge (Table 1), indicating the data was too limited to draw any conclusions. In two species (great tits and house sparrows) there was evidence that survival was higher closer to the city centre (Fig. 2a; Table 1, ΔAICc > 23.8). Here too, the change in survival associated with urbanization was substantial, with an increase of 19% in great tits and 12% in house sparrows living in city centres compared to those living 3.5km away from cities (Fig. S2). In general, both the distance from the city and IMP showed similar patterns, except for house sparrows and greenfinches (Fig. 2a & Fig S2). House sparrows had higher survival closer to the city centre, yet higher IMP was associated with lower survival (Fig. 2a & S2). Survival of greenfinches showed the opposite pattern with survival decreasing closer to the city centre, but increasing with higher IMP (Fig. 2a & S2), although the latter association was much weaker and did not receive model support (Table 1, ΔAICc = +1.7). Finally, modelling the urbanization predictors as quadratic effects on survival improved the model fit for several species (Table 1): the positive association between survival and distance to the city did not improve further once reaching the city border in great tits and to some extent also in blue tits (ΔAICc = 0.9; Fig. S2), whereas the negative association between IMP and survival was particularly apparent at high IMP for house sparrows and to some extent in blackcaps (*Sylvia atricapilla*, ΔAICc = 1.7; Fig S2).

**Fig. 2.**
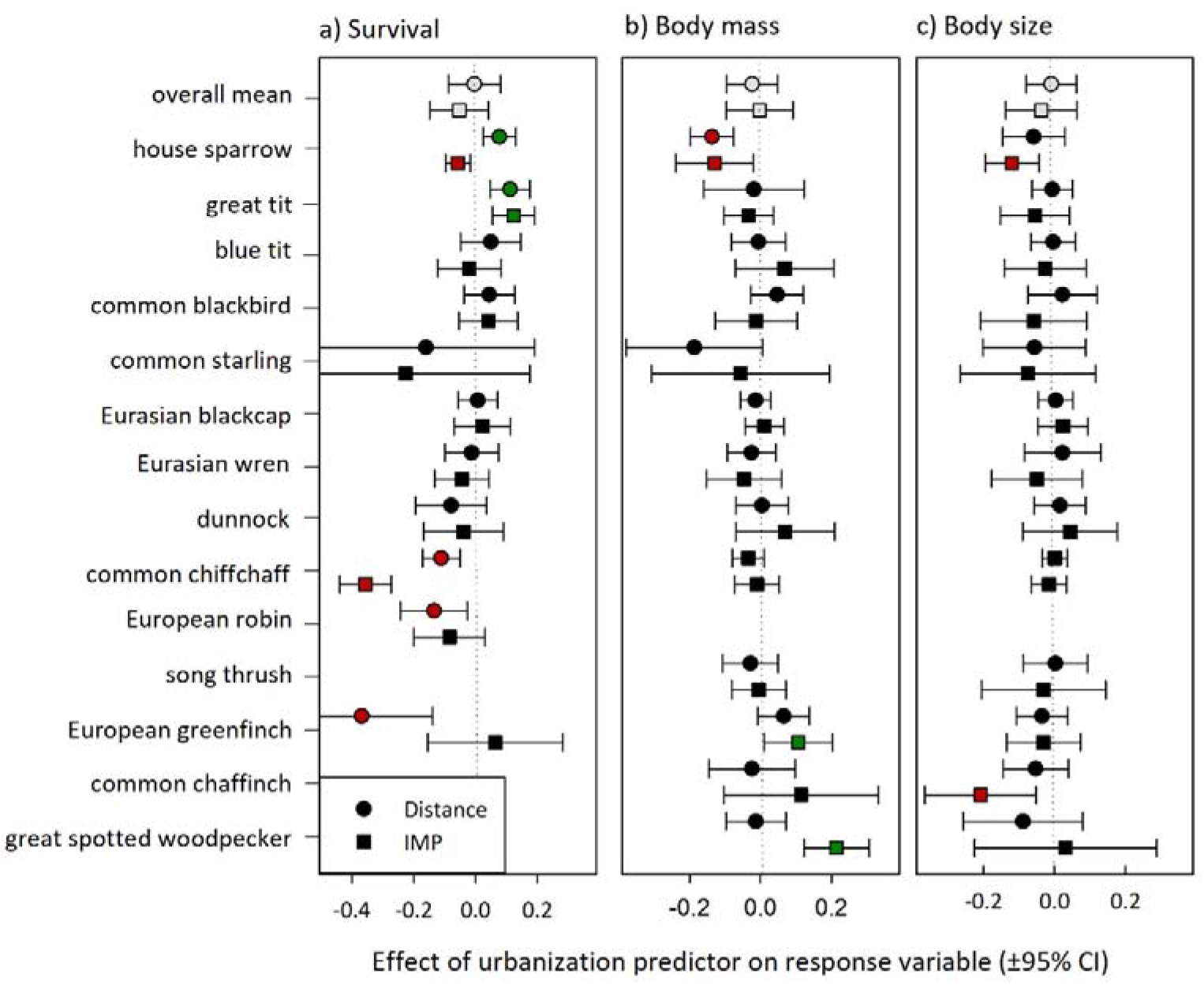
Standardized effects sizes (±95% CI) of the distance from the city border (distance) and the percentage of impervious surface (IMP) on a) apparent annual adult survival, b) body mass and c) body size (wing length). Positive effects are shown in green whereas negative effects are shown in red. Effect sizes for survival are on logit scale.

**Table 1.**
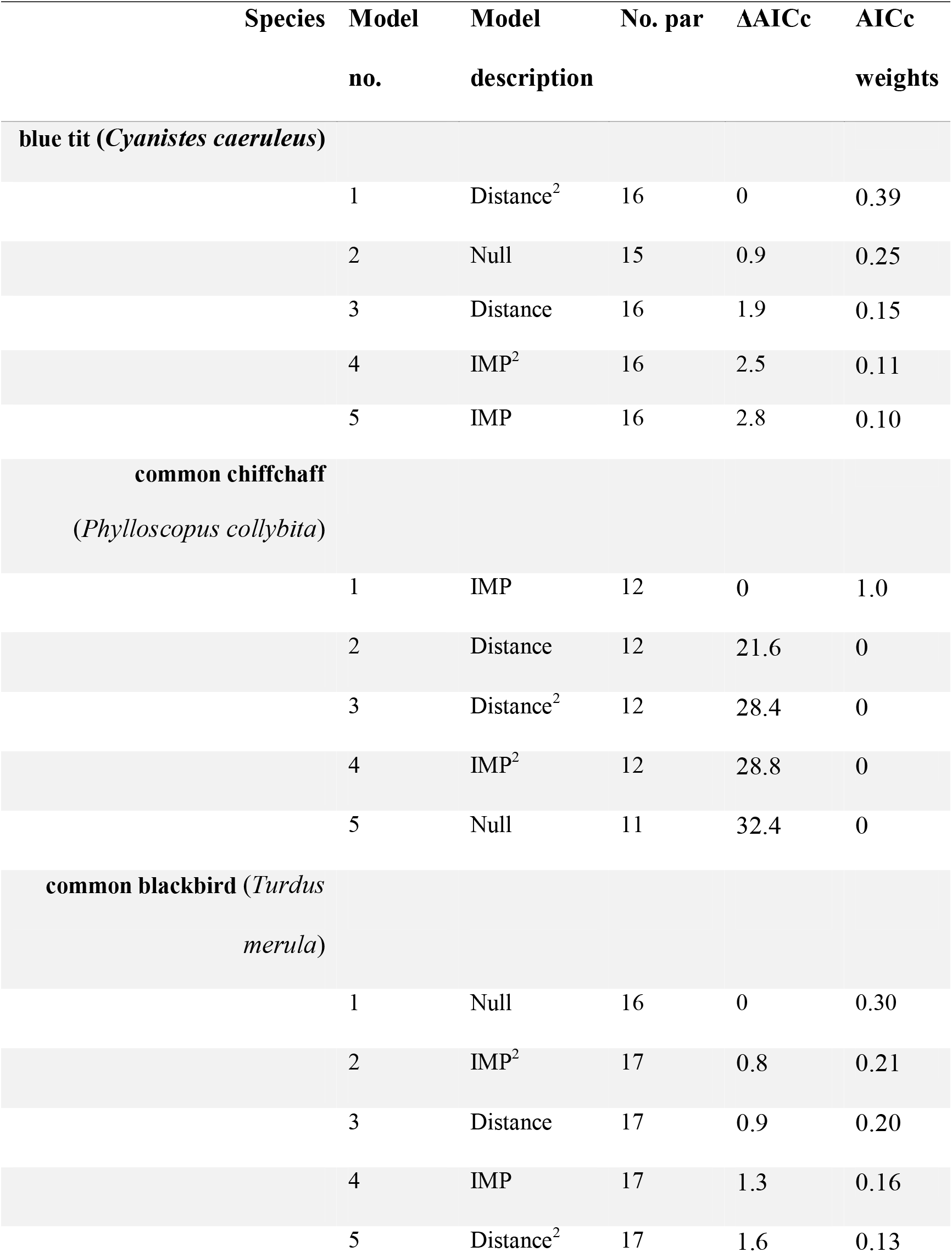
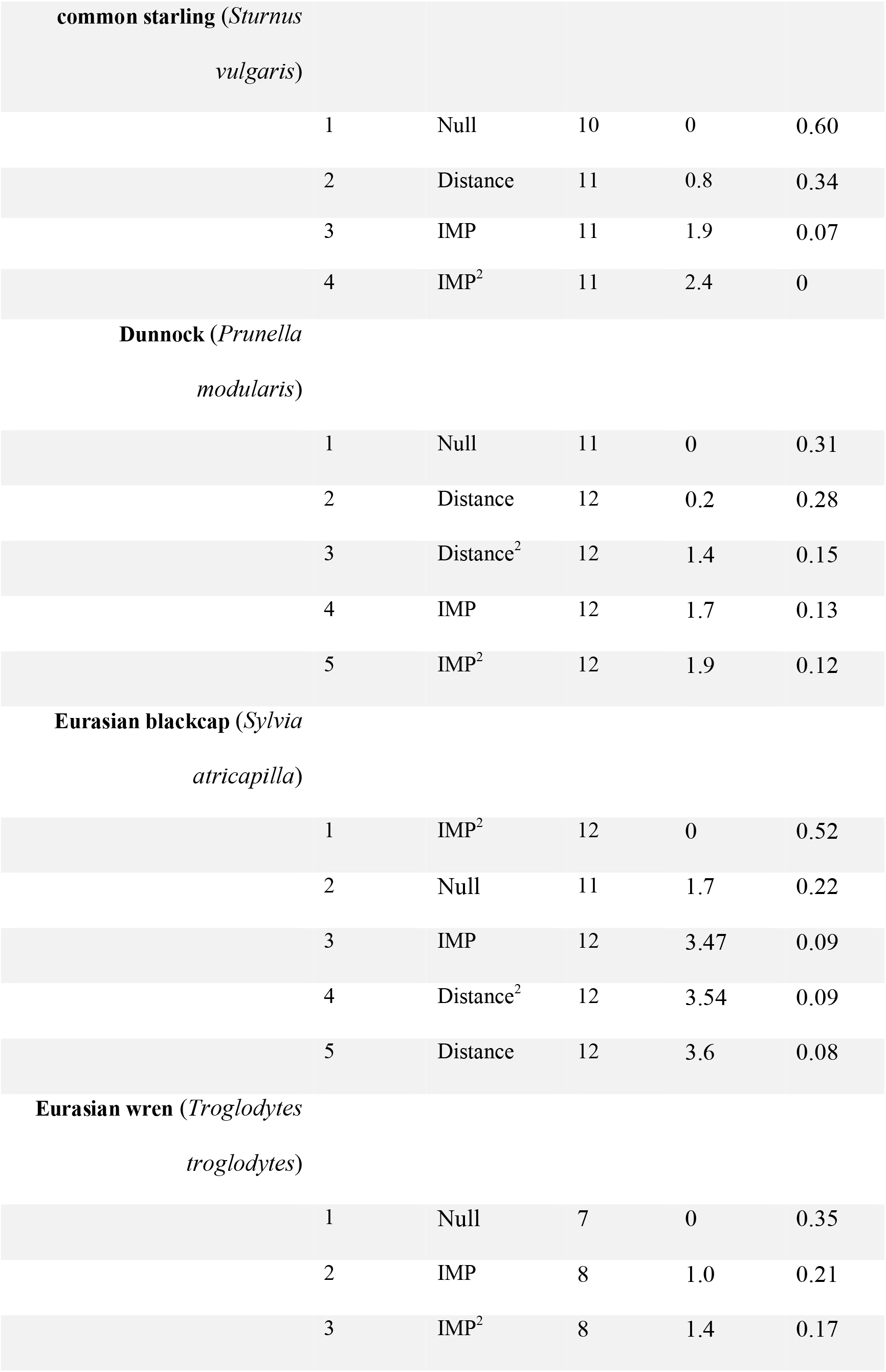
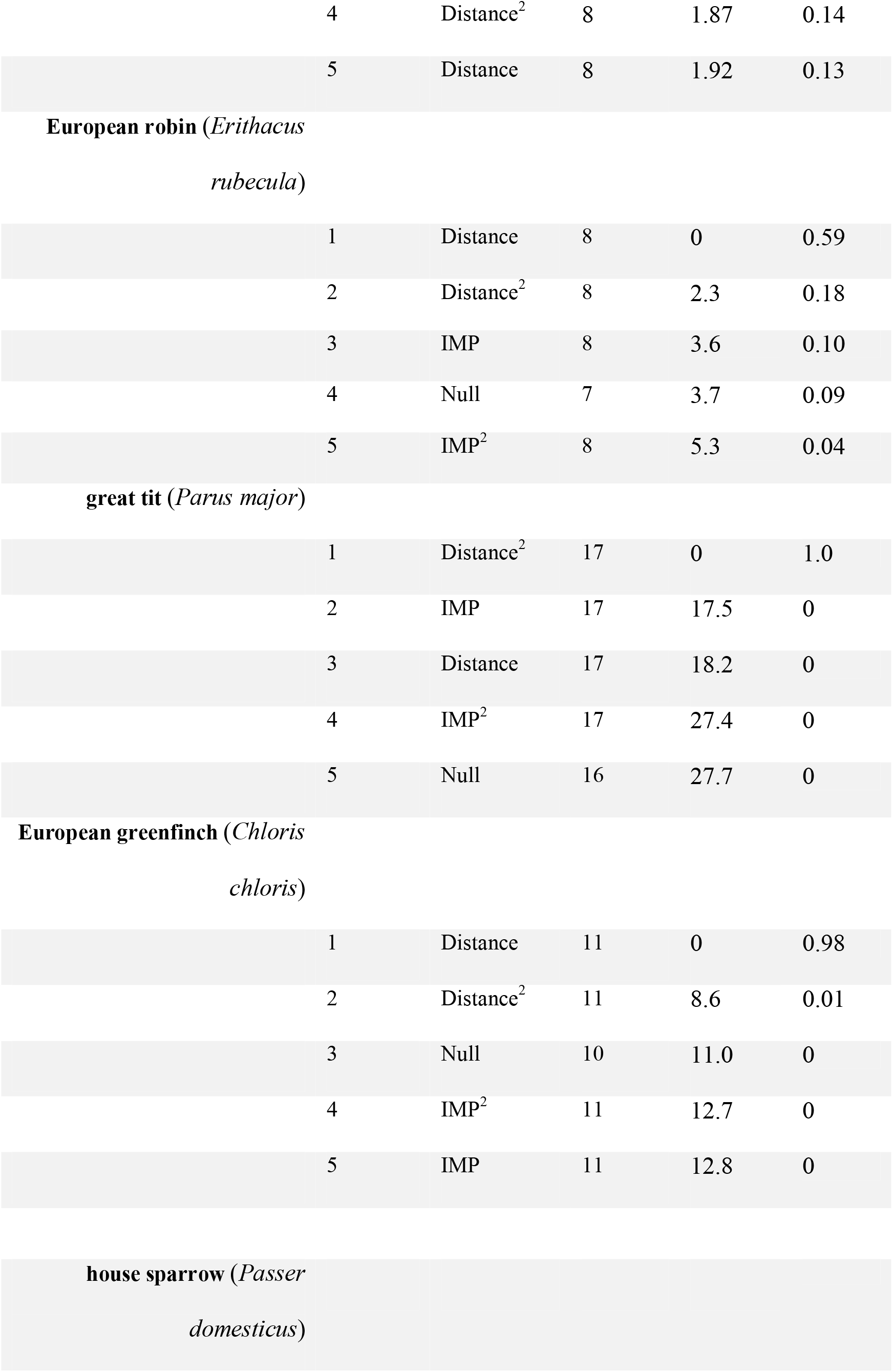
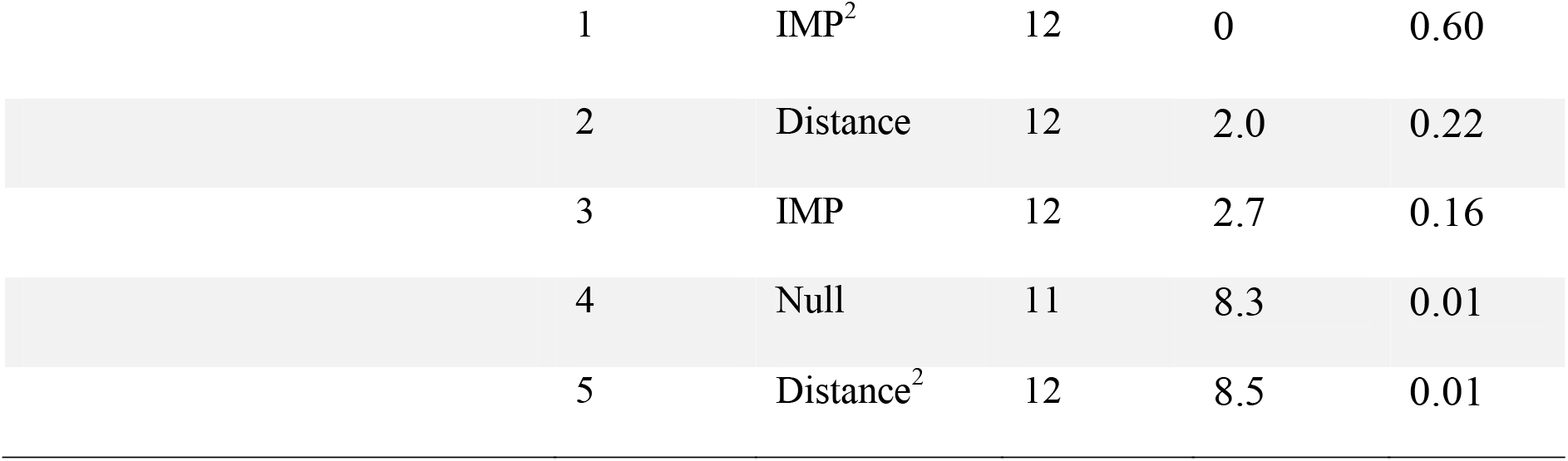
Summary of model selection statistics examining the effects of distance from the city border and percentage impervious surface (IMP) on apparent annual adult survival of 11 avian species. Models are ranked according to their ΔAICc.

### Relative body mass

Results on the structural equation models showed that in contrast to our prediction that urbanization has a negative effect on relative body mass, there was little evidence for such a general pattern: the average effect sizes of both urbanization predictors were only slightly negative with 95% CI’s that overlap zero (Fig. 2b). Interestingly, house sparrows were the only species where increased urbanization (both distance and IMP) was significantly associated with lower relative body mass (Fig 2b; Table S2). In contrast, greenfinches and great spotted woodpeckers had significantly higher relative body mass with higher levels of IMP (Fig. 2e, Table S2). Although a large proportion of the conditional (through random and fixed effects) variance in relative body mass was accounted for in the models (26-88%, Table S2), the amount of marginal variance in relative body mass that could be attributed to effects of urbanization was low at 3% in great spotted woodpeckers, 2% in house sparrows, and only <1% in greenfinches.

### Body size

Results on the structural equation models for body size (wing length) showed that in contrast to our prediction there was little support for an overall negative effect of urbanization on body size (Fig. 2c). Most effects were weak, and the majority of CI’s overlapped zero with only the effects of IMP for house sparrows and common chaffinch being significantly different from zero (Fig. 2c, Table S2). Similarly to the effects on mass, a large proportion of the conditional variance in body size was accounted for in the models (74-92%, Table S2), but the amount of marginal variance in body size explained through urbanization was very low at 2% in house sparrows and 1% in common chaffinches. Finally, there was no evidence that effects of urbanization on mass and size were mediated or traded-off against survival, as there was no significant correlation between effect sizes of survival and mass or size (*r*<0.25, *P*>0.48, df=8; Fig. 2).

## Discussion

Urbanization currently forms the most drastic change to the environment and since the expansion of urban land use is predicted to accelerate even further it will be crucial for conservation management to understand the consequences this has on wildlife populations. In addition to an individual’s ability to survive, increasing temperatures and changes in food availability associated with urbanization have also been predicted to result in variation in body mass and size. However, it is unclear how general such effects are given the largely mixed evidence from studies mostly focussing on single species, from few sites, over short timeframes. To resolve this important question, comparative studies on multiple species followed over large spatiotemporal scales are needed. Here, we investigated effects of urbanization on survival and biometry in common bird species collected over an 8-year period in a nationwide capture-mark-recapture study. Results showed strong variation among species. Although the overall association between urbanization and survival was slightly negative, there was strong support for lower survival in three and higher survival in two of the 11 species examined. Effects of urbanization on relative mass or size were detected in four of 13 species examined, but these also varied among species and their effects were very weak.

### Effects of urbanization on survival

Our results contrast with a recent meta-analysis, which found a positive association between survival and urbanization across avian species (Sepp et al., 2018). Such large-scale generalizations may not be meaningful when species show so much variation in their ability to cope with urbanization (Isaksson, 2018). Interestingly, our results show that two of the species showing reduced survival, the chiffchaff and European robin, are insectivorous, whereas the species that benefitted, great tits and house sparrows, are considered generalists. Our findings thereby corroborate the results from a multi-species study from the north-eastern USA, where particularly generalist species had higher survival with increased urbanization (Evans et al., 2015). Yet, given the heterogeneity in species survival responses, care should be taken with predictions based on species’ degree of specialism only. For example, the common starling is also typically considered a generalist species, but we found that its survival decreased with increased urbanization (although non-significantly so). Furthermore, even closely related species can vary in their response to urbanization. For example, whereas great tits had higher survival in areas with higher IMP, such a pattern was not detected for blue tits. Nevertheless, our results are largely in line with the consensus that generalists as opposed to specialists can thrive in the city (Møller, 2009; Marzluff, 2017; Isaksson, 2018). Given that avian reproductive success is generally lower in urban areas (even for generalists; Chamberlain et al., 2009; Sepp et al., 2018; Seress et al., 2020), the increased survival likely plays an important role in the success of these species.

These findings suggest that conservation actions should prioritize survival of insectivorous species in urban areas. For example, through increasing the availability of invertebrate prey by increasing green space like parks and the conversion of paved gardens to more natural habitat. However, it has been argued (Shochat et al., 2004; Evans et al., 2015) that lower survival due to urbanization mainly results from top-down processes like higher rates of predation (Bonnington, Gaston, & Evans, 2013; but see Fischer et al., 2012), or collisions with manmade objects (Loss et al., 2019), whereas higher survival results from bottom-up processes like higher resource availability (Shochat et al., 2004). This may suggest that for most species in our dataset top-down processes are relatively more important than bottom-up processes and that the predicted higher food availability cannot compensate for the negative top-down processes. Alternatively, the lower availability of natural food, such as insects (Jones & Leather, 2012; New, 2015; Seress et al., 2018) may mean that wildlife is more reliant on human derived food that may not cover the nutritional requirements (Shochat et al., 2004). Given that specifically insectivorous species suffered survival reductions, this is actually more likely.

Our results between the different measures of urbanization (IMP and distance to city border) largely corresponded, except for house sparrows which benefitted from living closer to and within cities, but surprisingly had lower survival in areas with more impervious surface. A possible explanation is that while house sparrows are opportunistic eaters, insects form an important part of their diet, both in urban and rural environments (Gavett & Wakeley, 1986). This suggests that the presence of suitable habitat to forage for insects might even be important for generalist species in urban areas. This idea is corroborated by the finding that the lower reproductive success of urban house sparrows in the UK could be (partly) explained by low aphid densities (Peach et al., 2008). The house sparrow, one of the best known urban dwellers, has gone through a 50% population reduction in western Europe since the 1980’s (Burns et al., 2021), and may thus also benefit from conservation actions focussing on increasing invertebrate prey in urban areas.

The strongest decline in survival with increasing urbanization was detected in the European greenfinch, a species which has been known to suffer from the highly infectious and fatal finch Trichomonosis disease caused by the parasite *Trichomonas gallinae* (Rijks et al., 2019). The extremely low survival of greenfinches in urban areas could largely be due to this disease, the spread of which is likely facilitated through an increased frequency of intra-specific interactions at feeding stations in gardens (Lawson et al., 2018). High resource availability through bird feeding may thus also result in negative effects on survival, although the role of urban bird feeding in disease systems is not yet well understood (Galbraith et al., 2017; Reynolds et al., 2017; Jones, 2018).

Capture-mark-recapture studies in open populations typically suffer from the problem that patterns in survival may be confounded with patterns in dispersal, because individuals permanently emigrating from the capture sites will be assumed dead. Movement of breeding adults (i.e. breeding dispersal) remains one of the least understood processes driving population dynamics (Greenwood & Harvey, 1982), not least because of the difficulties in studying such behaviour. In our study we attempted to reduce bias in survival estimates due to breeding dispersal by focussing on adult survival only (i.e. avoiding the natal dispersal phase); only using recaptures from the breeding season (i.e. from residents on their breeding grounds); and by accounting for transients (i.e. excluding individuals that were caught only once from contributing to the survival estimate) (Pradel et al., 1997, Johnston et al., 2016). Despite these measures, we cannot exclude the possibility that the observed survival patterns are driven by differences in breeding dispersal. As far as we are aware, only a single study examined such patterns, and found no evidence for differences in breeding dispersal of six songbird species among urban and natural landscapes in Washington, USA (Marzluff et al., 2016). The further development of tracking techniques will enable large scale studies on how patterns of dispersal vary along the urban gradient.

### Effects of urbanization on mass and size

Effects of urbanization on relative mass was generally weak, but nonetheless showed a few significant patterns. For example, despite their higher survival, house sparrows had lower relative body mass with shorter distance to the city. This may suggest that they can afford to carry less reserves (Shochat, 2004), an idea that is supported by recent experimental work which showed that supplementary fed urban birds had lower body condition than non-supplementary fed individuals (Demeyrier et al., 2017).

Another prediction from this ‘credit-card hypothesis’ is that the high food availability leads to a high proportion of weak competitors in urban environments although evidence for this is ambiguous (Shochat, 2004; Bókony, Kulcsár, & Liker, 2010) our findings that house sparrow survival is actually higher in urban areas does support this idea. In contrast to house sparrows, woodpeckers’ and greenfinches’ relative body mass increased closer to city centres. This may suggest that these species benefit from the food available in urban areas and do accumulate body reserves. Alternatively, the high mortality of greenfinches in urban areas may result in selective disappearance of individuals with lower relative body mass (Nussey et al., 2011).

Our study provided little evidence for consistent reductions in body size with increased urbanization, which was only found in house sparrows and chaffinches. Size reductions could be expected based on the lower food quality or from increased temperatures favouring smaller body size (Gardner et al., 2018, 2019). In nearby Belgium, urban-heat-island effects have been shown to drive invertebrate diversity towards smaller species (Merckx et al., 2018). Although the latter study also showed that filtering for smaller species can be over-ruled by filtering for larger species when there is positive covariation between size and dispersal. A similar process could occur within species and this could potentially explain the absence of an urbanization-size association in many of our studied species, where being larger may facilitate movement through fragmented urban areas.

## Conclusion

In conclusion, using data of 14 species collected over 8 years in >200 locations along a gradient of urbanization throughout the Netherlands showed strong variation in species responses. Effects of urbanization on body mass and size were very weak and showed that mass and size cannot be used as reliable quick indicators of species’ survival response to urbanization. The heterogeneity in responses means that care should be taken when predicting the effects of ongoing urbanization on communities, because even closely related species can show different responses. Nevertheless, our findings largely support the idea that generalists manage to benefit, whereas more specialist insectivorous species suffered from reduced survival. The increased survival of generalists likely plays the crucial role in the success of these species. Eventually, this is expected to lead to shifts in community composition through time and space and cause loss of biodiversity with ongoing urbanization. A conservation management objective to maintain avian biodiversity in urban areas should therefore focus on increasing invertebrate prey to promote survival of insectivorous species.

## Acknowledgements

We thank all the citizen scientists for their commitment to the various projects and Martijn van de Pol for helpful comments on the manuscript. This project has received funding from the European Union’s Horizon 2020 research and innovation program under the Marie Sklodowska-Curie grant agreement No 795921 awarded to LB. The Dutch Birdlife partner Vogelbescherming generously supported the start of the *ring*-MUS project in 2011, as well as the contribution of LdV and HvdJ to this paper.

## Data accessibility

Upon acceptance data used in this paper will be made available in an online repository.

