## Supplementary material for "A country-wide examination of effects of urbanization on common birds": supllement

**Supporting Information**

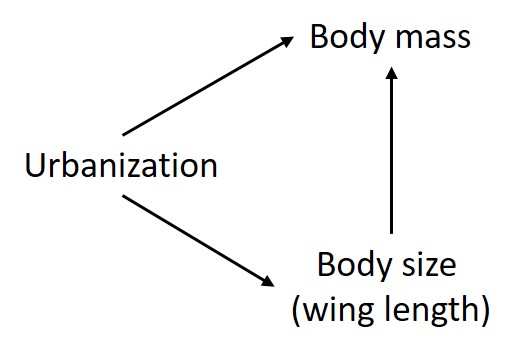

**Fig. S1** Path diagram showing the pathways in the structural equation model examining the effects of urbanization on body mass and size, while accounting for the effect of size on mass.

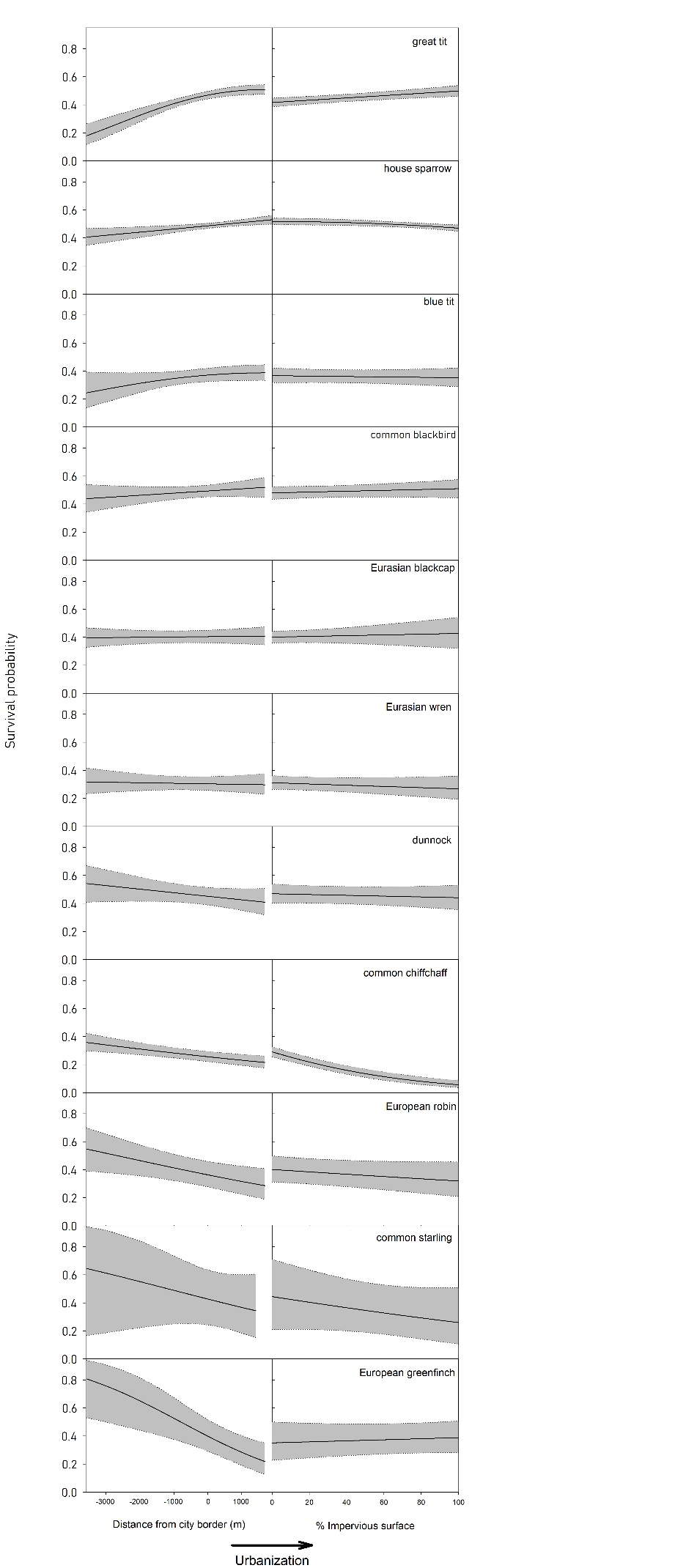

**Fig. S2.** Predicted survival probabilities (± 95CI) in relation to distance to the city border and percentage impervious surface for 11 bird species from the Netherlands. Estimates were derived from the models in Table 1 and include the linear effect of the urbanization predictors, except for great tit and blue tit (distance) and house sparrow and blackcap (IMP), where the results of the quadratic effects are shown (which received the strongest support: Table 1).

**Table S1.** Summary of sample sizes (number of individuals) and description of the null models used for parametrization of the survival and recapture parameters for each of the species used in the analyses.

| Species | N_biometry_ | N_survival_ | Null model survival | Null model recapture |
| --- | --- | --- | --- | --- |
| song thrush (*Turdus philomelos*) | 838 | NA | NA | NA |
| great spotted woodpecker (*Dendrocopos major*) | 323 | NA | NA | NA |
| European robin (*Erithacus rubecula*) | NA | 4,306 | Age | Age+RingAge |
| common blackbird (*Turdus merula*) | 2,363 | 4,772 | Age | Age+RingAge+Project+RingType+RingAge*Project+Age*Project |
| Eurasian blackcap (*Sylvia atricapilla*) | 5,360 | 11,884 | Age | Age+RingAge+Project+RingAge*Project+Age*Project |
| common starling (*Sturnus vulgaris*) | 296 | 2,311 | Age | Age+RingAge+Project+RingType |
| dunnock (*Prunella modularis*) | 1,254 | 2,682 | Age | Age+RingAge+Project+RingAge*Project+Age*Project |
| common chiffchaff (*Phylloscopus collybita*) | 3,241 | 13,292 | Age | Age+RingAge+Project+RingAge*Project+Age*Project |
| Eurasian wren (*Troglodytes troglodytes*) | 1,666 | 3,908 | Age | Age+RingAge |
| European greenfinch (*Chloris chloris*) | 1,725 | 3,018 | Age | Age+RingAge+Project+ RingType |
| common chaffinch (*Fringilla coelebs*) | 866 | NA | NA | NA |
| house sparrow (*Passer domesticus*) | 3,128 | 14,761 | Age | Age+RingAge+RingType+RingAge*RingType+Age*RingType |
| great tit (*Parus major*) | 3,051 | 14,582 | Age | Age+RingAge+Project+RingType+RingAge*Project+Age*Project |
| blue tit (*Cyanistes caeruleus*) | 1,431 | 8,913 | Age | Age+RingAge+Project+RingAge*Project+Age*Project |

**Table S2.** Results from structural equation modelling showing effects sizes (± S.E.) for the effects of distance to the city edge and percentage impervious surface on body weight and wing length of 13 bird species. R^2^ is the conditional value obtained from the null model (without the urbanization predictors).

| Species | Trait | Distance | % IMP | Distance^2^ | IMP^2^ | R_Null_^2^ |
| --- | --- | --- | --- | --- | --- | --- |
| blue tit | weight | -0.006 ± 0.039 | 0.068 ± 0.071 | -0.006±0.037 | 0.075 ± 0.054 | 0.54 |
|  | wing | -0.004 ± 0.032 | -0.027 ± 0.059 | 0.007 ± 0.031 | -0.033 ± 0.045 | 0.81 |
| common chiffchaff | weight | -0.035 ± 0.023 | -0.01 ± 0.032 | -0.028 ± 0.023 | 0.008 ± 0.030 | 0.26 |
|  | wing | 0.000 ± 0.018 | -0.016 ± 0.025 | -0.006 ± 0.018 | -0.032 ± 0.022 | 0.88 |
| common blackbird | weight | 0.047 ± 0.038 | -0.013 ± 0.059 | 0.042 ± 0.037 | -0.004 ± 0.054 | 0.51 |
|  | wing | 0.022 ± 0.05 | -0.059 ± 0.076 | 0.025 ± 0.047 | -0.048 ± 0.063 | 0.77 |
| common chaffinch | weight | -0.024 ± 0.062 | 0.114 ± 0.111 | -0.041 ± 0.042 | 0.096 ± 0.072 | 0.69 |
|  | wing | -0.054 ± 0.047 | **-0.209 ± 0.08** | -0.017 ± 0.032 | -0.0161 ± 0.054 | 0.88 |
| common starling | weight | -0.187 ± 0.098 | -0.057± 0.128 | -0.189 ± 0.089 | -0.105 ± 0.126 | 0.88 |
|  | wing | -0.057 ± 0.074 | -0.076 ± 0.097 | -0.062 ± 0.069 | -0.077 ± 0.095 | 0.92 |
| dunnock | weight | 0.004 ± 0.038 | 0.069 ± 0.071 | 0.005 ± 0.036 | 0.031 ± 0.055 | 0.55 |
|  | wing | 0.015 ± 0.037 | 0.044 ± 0.068 | 0.008 ± 0.035 | -0.077 ± 0.095 | 0.85 |
| Eurasian blackcap | weight | -0.014 ± 0.022 | 0.011 ± 0.028 | 0.018 ± 0.021 | 0.009 ± 0.028 | 0.48 |
|  | wing | 0.003 ± 0.025 | 0.023 ± 0.036 | 0.002 ± 0.024 | 0.003 ± 0.031 | 0.75 |
| Eurasian wren | weight | -0.026 ± 0.035 | -0.047 ± 0.054 | -0.021 ± 0.035 | -0.027 ± 0.043 | 0.47 |
|  | wing | 0.022 ± 0.055 | -0.05 ± 0.065 | 0.017 ± 0.058 | -0.027 ± 0.043 | 0.74 |
| European greenfinch | weight | 0.065 ± 0.037 | **0.106 ± 0.049** | 0.055 ± 0.033 | **0.124 ± 0.044** | 0.50 |
|  | wing | -0.036 ± 0.037 | -0.032 ± 0.053 | -0.044 ± 0.032 | -0.074 ± 0.049 | 0.78 |
| great tit | weight | -0.019 ± 0.031 | -0.034 ± 0.047 | 0.003 ± 0.030 | -0.022 ± 0.038 | 0.47 |
|  | wing | -0.007 ± 0.029 | -0.056 ± 0.050 | 0.009 ± 0.029 | -0.060 ± 0.036 | 0.81 |
| house sparrow | weight | **-0.137 ± 0.04** | **-0.13± 0.036** | -0.171 ± 0.041 | -0.095 ± 0.035 | 0.58 |
|  | wing | -0.059 ± 0.045 | **-0.12 ± 0.039** | -0.069 ± 0.046 | -0.125 ± 0.038 | 0.89 |
| great spotted woodpecker | weight | -0.013 ± 0.072 | **0.214± 0.100** | -0.026 ± 0.070 | **0.191 ± 0.091** | 0.45 |
|  | wing | -0.089 ± 0.086 | 0.031 ± 0.131 | -0.058 ± 0.085 | 0.108 ± 0.113 | 0.78 |
| song thrush | weight | -0.029 ± 0.035 | -0.005± 0.056 | -0.031 ± 0.034 | 0.001 ± 0.045 | 0.64 |
|  | wing | 0.002± 0.046 | 0.031 ± 0.089 | 0.010 ± 0.045 | -0.053 ± 0.062 | 0.85 |
